# Classifying bladder cancer subtypes

**DOI:** 10.1101/250258

**Authors:** Cihan Kaya, Nicolas Arcenio Pabon

**Affiliations:** Department of Computational and Systems Biology, University of Pittsburgh, Pittsburgh, PA 15213

## Abstract

Urothelial carcinoma of the bladder is is estimated to have killed over 16,000 people in the United States in 2016. Like breast cancer, bladder cancer is a heterogeneous disease, and characterization of it’s various subtypes can be useful for forecasting prognosis and treatment efficacy. According to The Cancer Genome Atlas (TCGA) project, the mRNA expression profiles of bladder tumours can be used to cluster the tumors into four different categories: I - Papillary-like, II ‐Luminal A, III - basal/squamous-like, and IV - other (similar to III). However it is not clear whether these mRNA expression based clusters correlate with other molecular and genetic features of the tumor cells. In other words, do differences in mRNA expression profile contain the same information as differences in protein expression, micro RNA (miRNA) expression, copy number variation and somatic mutation data. We tried to recreate mRNA based bladder tumor clusters from other multi-omic data for 328 bladder cancer tumor samples using a special deep and wide belief network composed of restricted Boltzmann machines and a multilayer perceptron. For 10-fold cross validation, we got 79% average test accuracy which implies that that differences in mRNA expression between bladder tumor cells can be reliably, though not perfectly, inferred from different molecular and genetic features of the tumors.

## 1 Introduction

Urothelial carcinoma of the bladder is diagnosed in 380,000 people and causes 150,000 deaths per year worldwide [1]. Although it has long been recognized that bladder cancer is heterogeneous and comprises multiple distinct disease entities with different molecular and genetic features and clinical outcome [2], comprehensive studies attempting to formally classify different bladder cancer subtypes have been lacking in comparison to other cancers such as breast cancer, head and neck cancer, and lung cancer. Since identifying cancer subtype can yield valuable information about likely tumor progression pathways and treatment resistance, investigating methods to formally categorize bladder cancers holds promise for reducing the global impact of the disease.

The most comprehensive analysis of the molecular and genetic landscape of of bladder cancer that has appeared so far was carried out by The Cancer Genome Atlas (TCGA) project [3,4]. The analysis includes demographic, clinical, and pathological data for 129 tumor samples as well as whole exome sequencing, RNA sequencing, miRNA profiling, protein expression, and several genetic markers such as somatic mutations, chromosome rearrangements, viral integrations, and copy number variations. The study used a bootstrapped ensemble clustering algorithm to divide the tumor samples into four distinct categories (Clusters 1-4) using mRNA expression profiles from 2708 landmark genes (Figure 1).

**Figure 1:**
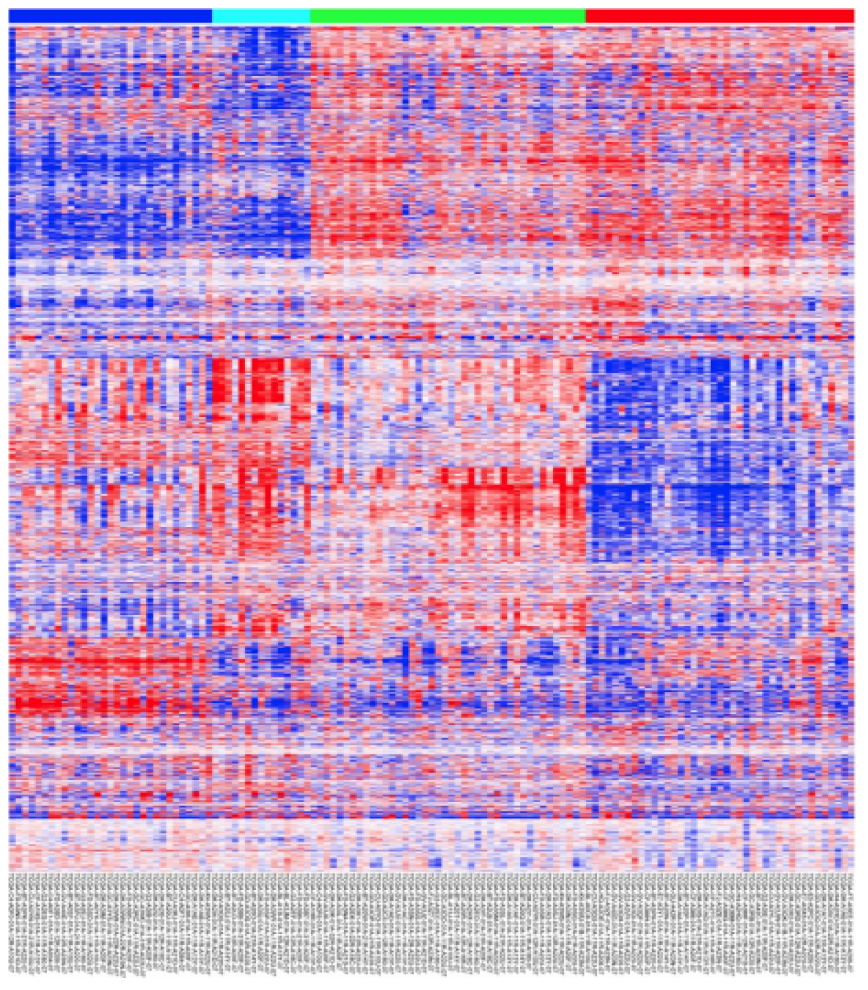
RNA expression - Illumina HiSeq for 2708 variable genes in 129 TCGA bladder cancer (BLCA) samples. The four different clustered are indicated by the different colors in the top bar.

Among other distinctions between the clusters, the authors found that Cluster 1 was enriched in tumors with papilary morphology, and, along with Cluster 2, showed features similar to those of luminal A breast cancer, with high mRNA and protein expression of luminal breast differentiation markers. The mRNA signature or bladder cancer Clusters 3 and 4 were found to be similar to those of basal-like breast cancers, as well as squamous cell cancers of the head and neck and lung (Figure 2). Various correlations of the clustering scheme with differential protein and miRNA expression levels as well as gene mutation and amplification were also described (Figure 3).

**Figure 2:**
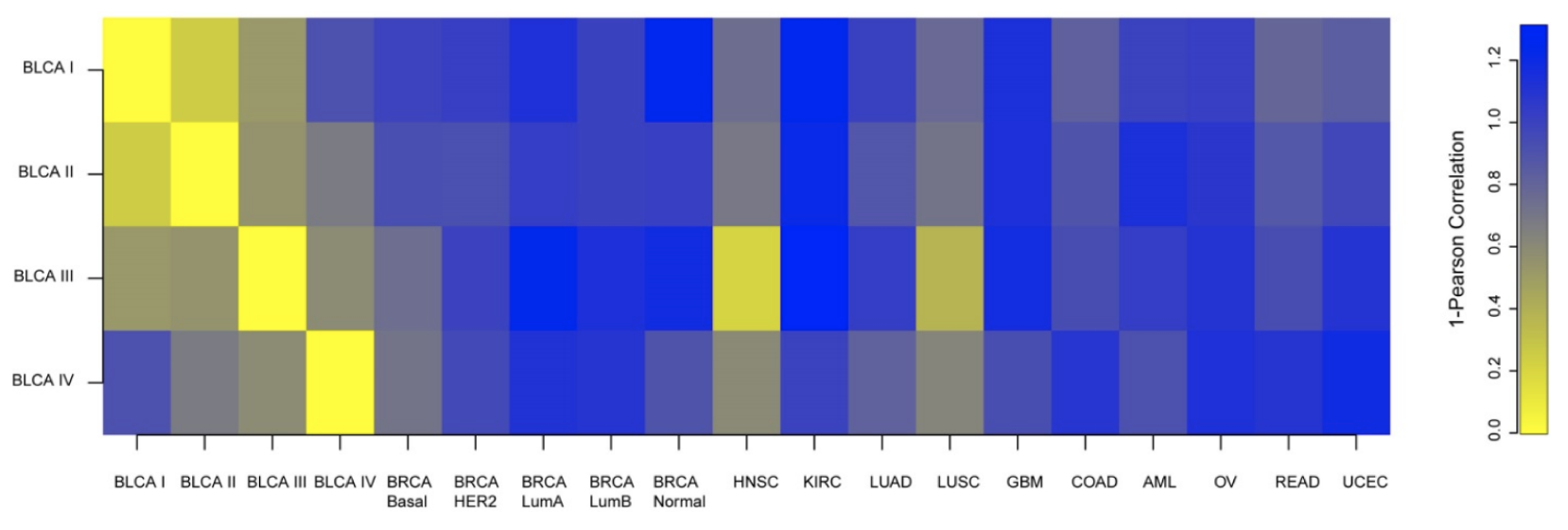
RNA bladder tumor subtypes and correlation to other TCGA tumor types

**Figure 3:**
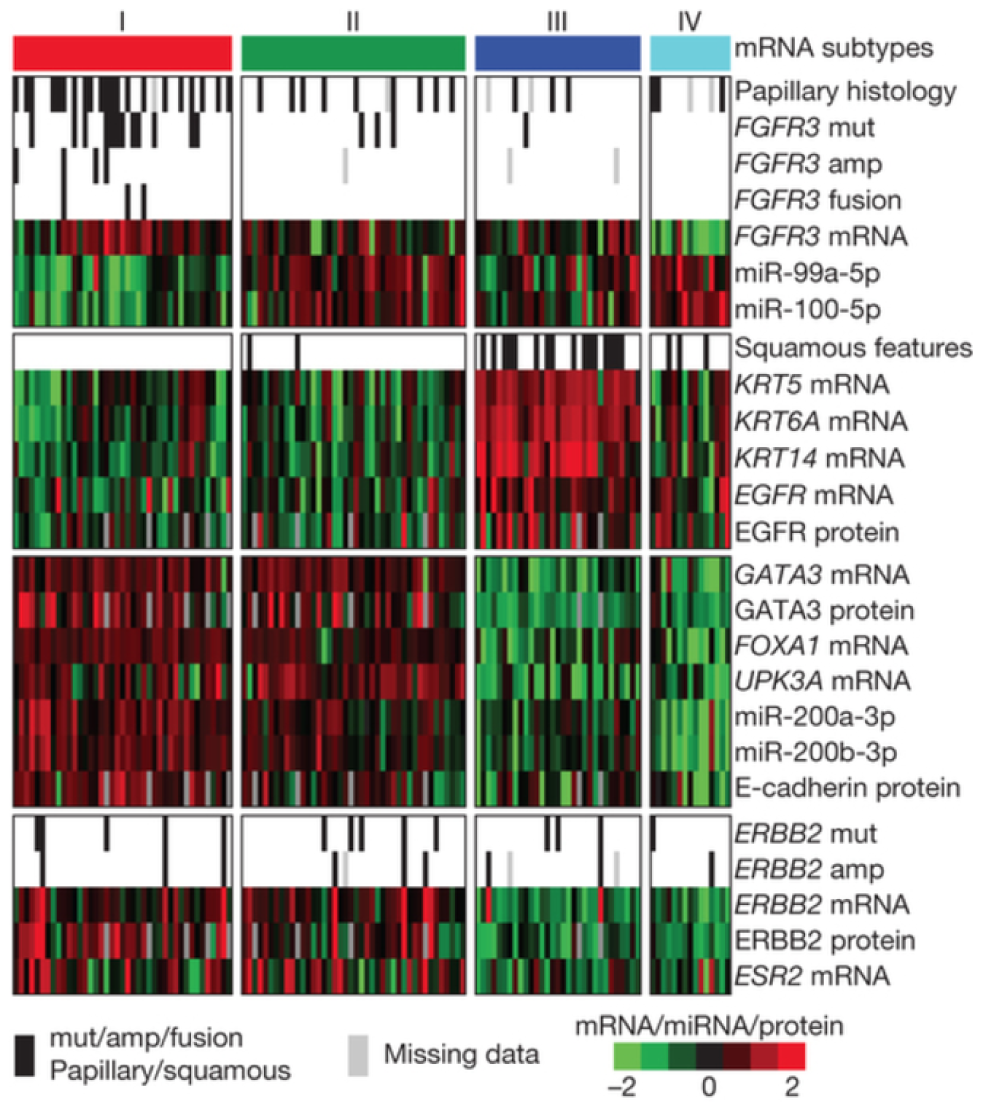
Expression characteristics of bladder cancer subtypes

Although the authors of the TCGA study mentions a number of specific non-mRNA tumor features (gene mutations & amplifications, miRNA & protein expression, etc.) that seem to align well with with the mRNA based clustering scheme (Figure 3), a more robust analysis comparing the different feature landscapes is needed. While it seems possible that mRNA expression level may serve as a reasonable proxy for cellular features such as protein expression level, it is less clear how changes in mRNA expression might correlate with copy number variance, somatic mutations, or miRNA expression levels.

Here, we attempt to answer one part of this complex and multi-faceted question, namely: is mRNA expression level contain unique information about the tumors, or can the same clusters be reasonably recreated using other molecular and genetic features. To this end we develop a deep belief neural network architecture that we train to classify 328 bladder tumor samples into the four categories from [3] without using the mRNA expression data that was used to perform the original clustering. Instead we attempt to classify these tumors using protein expression more specifically reverse phase protein array (RPPA), miRNA expression, somatic mutation, and copy number variation (CNV) data. Because of the high dimensionality of our input data, we use Restricted Boltzmann Machines to autoencode each data stream before passing them to a fully connected, feed forward neural network for classification. Our average classification accuracy for 10-fold cross validation is 79%, suggesting that the signal of the mRNA expression landscape can be reasonably reconstructed from alternative molecular and genetic features of the tumor cells.

## 2 Methods

### 2.1 Data Acquisition

We use UCSC Xena system [5] to access our download our desired tumor data from the TCGA [4]. Data is made available as text files that we downloaded and processed with Python and Scala. We download all available mRNA expression, RPPA, miRNA expression, somatic mutation, and CNV datasets for the TCGA Bladder Cancer (BLCA) cohort. Not all types of data are available for all tumors, and since we would like to use all data types for classification, we take as our training/validation set the intersection of all four datasets, which contains 328 tumor samples.

### 2.2 Assigning clusters to new tumors

The TCGA BLCA cohort contained 129 tumor samples at the time of the original study [3]. Now, it contains 328 samples, which include the original 129 tumors and others that have been profiled since the time of the study. Since we would like to use all available TCGA BLCA data to train/validate our model, we needed to assign cluster labels to the 199 new tumors. We took a k-means approach using, defining the cluster centers as the average of the mRNA expression vectors (d = 2708) of the already labelled tumors in each cluster from the original study. For the remaining 199 tumors, we extracted the relevant mRNA expression signatures from the mRNA dataset, determined the closest cluster center, and labelled each tumor sample with the cluster label. Data processing and tumor labelling was performed in Scala.

### 2.3 Data Normalization

Our original multi-omic data set, which included RPPA, CNV, somatic mutation, and miRNA expression data for each tumor (but not mRNA expression data, which was used exclusively for assigning labels to new clusters), was heterogeneous (different data types took on different values) and had high dimensionality of 68,265. We noticed that many features were uniform accross all, or almost all, tumor samples, meaning that they held no discriminatory power and would be uninformative in classification. Thus, in order to amplify the signal of our input data, and reduce the training overhead, we normalized each data channel to eliminate uninformative features and standardize the feature ranges. Collectively, normalization reduced the dimensionality of our data by almost half to 38,279.

#### 2.3.1 RPPA data normalization

The original RPPA data set contains 344 patients, each with 246 identifiers. Inspection of the data set revealed several missing values for some identifiers. We removed those identifiers to reduce total number of identifiers to 195. The remaining data was distributed with mean 0.02 and with variance 0.24. We normalized the data to fit between the -1 to 1 interval.

#### 2.3.2 miRNA data normalization

The original miRNA data set contains 429 patients, each with 2,221 identifiers. Inspection of the data revealed that many miRNA identifiers had multiple missing values for multiple patients. We removed those identifiers, reducing the dimensionality of the miRNA data to 1047. The remaining miRNA expression data is positive floating point data with mean 1.49 and variance 11.46. This data is normalized to fit between the -1 and 1 interval.

#### 2.3.3 CNV data normalization

The original CNV data set contains 408 patients, each with 24,777 genes that have copy number variation information. The mean value of the data is 0.03 and the variance is 0.16. In order to amplify the signal coming from important genes, we eliminated gene identifiers whose average variation was less than 0.1, reducing dimensionality to 16013. We then normalized values to fit between the -1 and 1 interval.

#### 2.3.4 Somatic mutation data normalization

The original somatic mutation data set contains 405 patients, each with 41,021 binary identifiers. The data set is 99.46% negative, making this a rare positive situation. We removed all genes with negative values for all patients, reducing dimensionality to 21,024. We changed all zero values to -1 so that the data fit between the -1 and 1 interval, as in the case of the other data types.

### 2.4 Network Architecture

Even after data normalization, the dimensionality of each tumor sample data point was over 38,000. Initial attempts to classify these points using a multilayer perceptron trained very slowly and we failed to achieve classification accuracy above 30%. At this point we looked for ways to amplify the signal to noise ratio in the data and arrived at the network architecture shown in Figure 4. Our general approach was to use autoencoders to reduce the dimensionality of our input data, and then feed the four data channels into a multilayer perceptron for classification.

**Figure 4:**
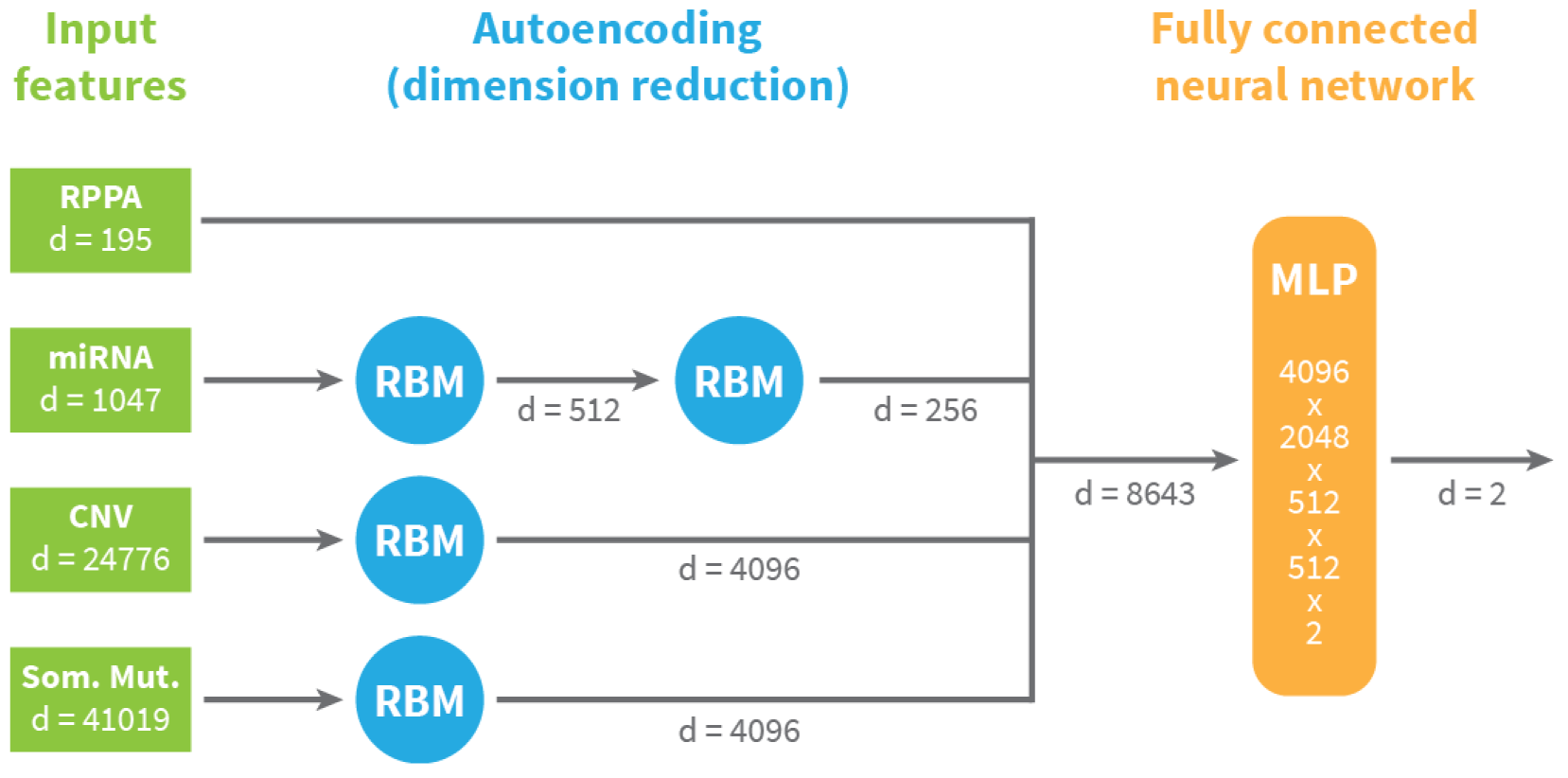
Deep belief network architecture of classifying bladder cancer tumor subtypes

#### 2.4.1 Autoencoding tumor data with Restricted Boltzmann Machines

For autoencoding we used Restricted Boltzmann Machines (RBMs) [6], generative stochastic artificial neural networks that learn probability distributions over their inputs. Briefly, an RBM is a bipartate graph of neurons with bidirectional edges between the visible and hidden layers. In our implementation, the visible units are the normalized tumor data (miRNA, RPPAs, etc…), which we assume are generated by a reduced set of hidden units that encode the tumor data. RBM training attempts to maximize the product of probabilities of the visible units in some training set by running a gradient descent procedure on the weight and offset values of the graph. Because training RBMs does not require back propagation we can train them much faster than a feed forward neural net and we can train them in parallel for each data channel. We trained our RBMs using the contrastive divergence algorithm [7] in TensorFlow [8] and attempted to minimize the reconstruction loss of the input data.

#### 2.4.2 Classifying tumors using multilayer perceptron

Autoencoding reduced the dimensionality of our tumor data to 8643, and this encoded data was fed into a feed forward neural network for classification. We tested various network architectures in a random search, using the Python scikit-learn [9] package for training, and found that a six layer fully connected network of dimension 8643 x 2048 x 1024 x 512 x 512 x 4 yielded the best classification accuracy in cross validation. L2 regularization was used during training to restrict model complexity and prevent overfitting.

## 3 Results

We partitioned our 328 tumor samples and performed 10-fold cross validation to try and classify each tumor in the validation set into the appropriate mRNA-based cluster.

### 3.1 RBM training results

Because of the low dimensionality of the RPPA data, we chose to bypass the autoencoding for this channel. We also found that stacking two RMBs in the miRNA channel produced lower reconstruction error than a single RBM while reducing the dimensionality by a factor of 2. This was not the case for the CNV and somatic mutation data channels; additional RBMs increased training time but did not improve reconstruction so we used only one RBM for these data types. The hyperparameters and final reconstruction loss for each of our RBMs is shown in Table 1.The change in the reconstruction loss per training epoch is shown for each RBM in Figure 5. Due to time constraints, the hyperparameters were not thoroughly optimized, and in theory reconstruction error might be improved even further through a hyperparameter grid search in future work.

**Table 1:**
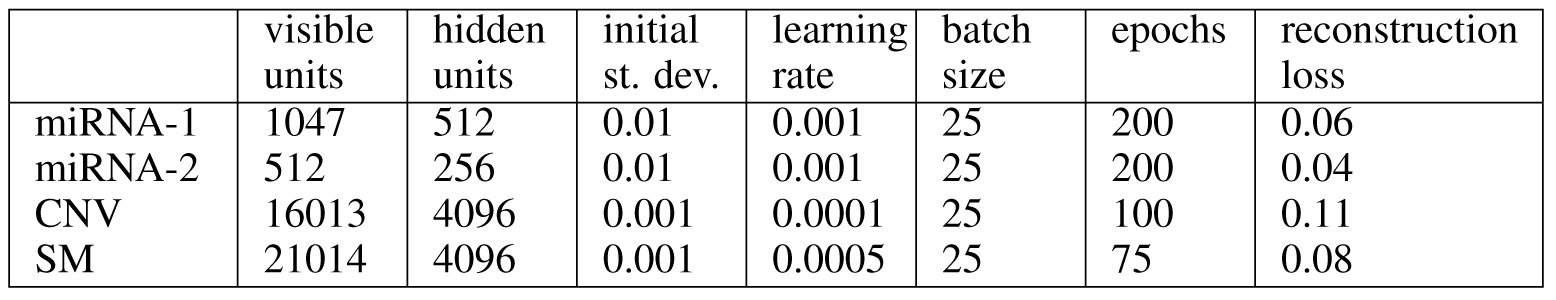
Hyperparameters and reconstruction loss for individual RBMs

|  | visible units | hidden units | initial st. dev. | learning rate | batch size | epochs | reconstruction loss |
| --- | --- | --- | --- | --- | --- | --- | --- |
| miRNA-1 | 1047 | 512 | 0.01 | 0.001 | 25 | 200 | 0.06 |
| miRNA-2 | 512 | 256 | 0.01 | 0.001 | 25 | 200 | 0.04 |
| CNV | 16013 | 4096 | 0.001 | 0.0001 | 25 | 100 | 0.11 |
| SM | 21014 | 4096 | 0.001 | 0.0005 | 25 | 75 | 0.08 |

**Figure 5:**
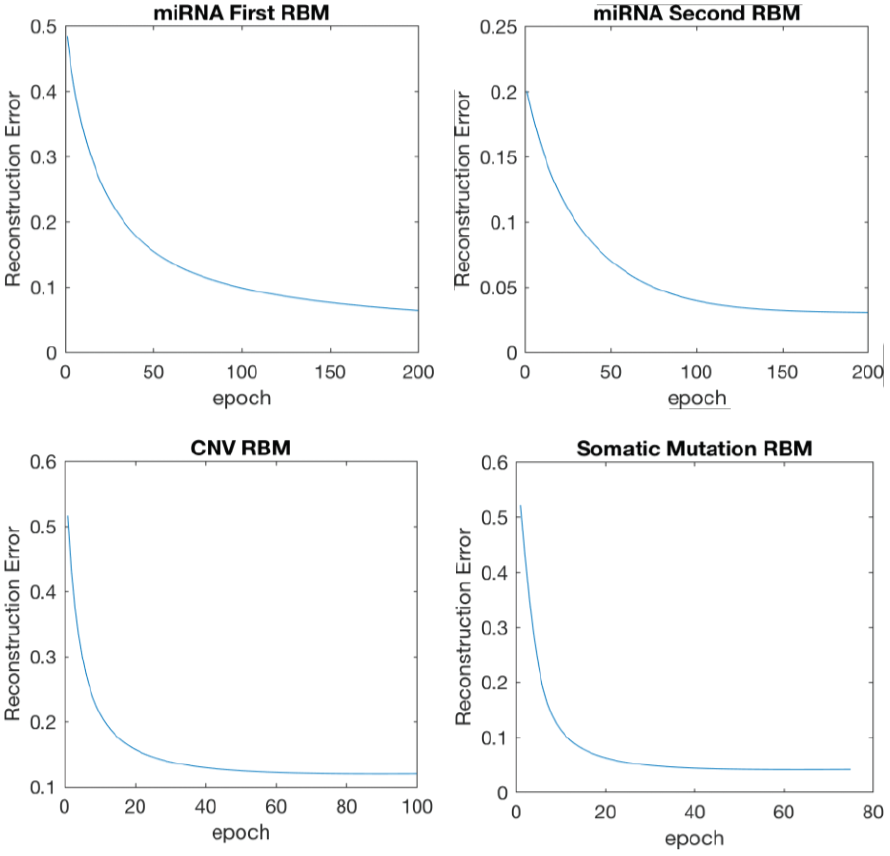
RBM reconstruction loss during training

### 3.2 Tumor classification results

We tested various multilayer perception architectures in a 10-fold cross validation procedure with an L2 regularization penalty of 0.001, learning rate of 0.0001, batch size of 25, and we sampled the initial model parameters from a truncated normal distribution with standard deviation 0.01. The 8643 x 2048 x 512 x 512 x 4 network architecture yielded the best performance of the networks that we tested, with an average classification accuracy of 35%. While this performance is better than random classification into the 4 Clusters, it is obviously not ideal.

We compared the predicted vs. true labels for our various tumor samples and found that the most frequent errors were misclassifications between Clusters 1 & 2, and between Clusters 3 & 4. In other words, our model was good at distinguishing Clusters 1 & 2 from 3 & 4, but not good at distinguishing 1 from 2, or 3 from 4. This was not a surprising result; the 4 Clusters defined in the original TCGA study [3] were grouped based on mRNA data. As can be seen in Figure 1, there is clear distinction in the expression patterns between the four Clusters. However, when the authors of [3] looked at other correlated expression characteristics (protein, mrNA, etc…), the boundary between Clusters 1 & 2 and between Clusters 3 & 4 became less clear (see Figure 3).

Thus we modified our tumor labels, combining Clusters 1 & 2 and Clusters 3 & 4, and we modified our network architecture to have two terminal nodes instead of four. This increased our average 10-fold cross validation accuracy to 79% after 100 epochs of training.

## 4 Discussion

We sought out to determine if the cellular mRNA profiles of the TCGA bladder cancer cohort contained unique information about the tumors that could not be determined from other molecular and genetic markers. To this end, we built a multi-channel deep belief neural network to classify the 328 bladder tumors in the TCGA into the clusters from the most comprehensive bladder cancer study [3]. We autoencoded our data using RBMs to reduce dimensionality and classified the encoded data using a 6 layer feed forward neural network. 10 fold cross validation of our final network architecture yielded 35% accuracy using the 4-Cluster scheme from the original TCGA study, and 79% accuracy when we combined Clusters 1 & 2 and 3 & 4.

First, we note that 79% accuracy of a binary classifier is far from perfect when we consider that the expected accuracy of a random classifier would be 50%. Due to time constraints on this work, we were not able to perform robust hyperparameter or model architecture searches, which might have improved classification accuracy, perhaps to a significant degree. If this were the case, we would discover that the signature of cellar mRNA profiles could was redundant; i.e. it’s information is reliably contained in the form of DNA copy number variations, somatic mutations, and protein & miRNA expression levels.

However, the degree to which model tweaking and parameter optimization might improve performance is unclear. There seems to be no justifiable reason to doubt that the cellular mRNA landscape contains some information that does not correlate with patterns in other molecular and genetic features, or perhaps that the correlation occurs on longer timescales than the experimental data acquisition window. Presumably if there is a short timescale correlation we should be able to detect it, perhaps by considering additional TCGA data in our model such as tumor phenotype, DNA methylation, and/or other metrics.

Ultimately, our results demonstrate the mRNA landscape of bladder cancer tumor cells is partially correlated with other types of multi-omic data. Given that current characterizations of bladder cancer subtypes are relatively young compared to other cancers, and are likely to evolve as new data and analyses emerge for this disease, for now our proposed methodology can serve to suggest subtype classes in case of missing mRNA expression data for new patients. These classifications could prove useful in suggesting further medical tests, determining treatment strategy, and forecasting disease progression.

